# The role of the innate immune system in shaping the dynamics of antimicrobial treatment

**DOI:** 10.64898/2026.04.09.717387

**Authors:** Teresa Gil-Gil, Brandon A. Berryhill, Christopher Witzany, Roland R. Regoes, Fernando Baquero, Bruce R. Levin

## Abstract

Contemporary antibiotic treatment almost solely considers the Minimum Inhibitory Concentration (MIC) when deciding to employ an antibiotic, often overlooking the critical role of host immunity. Using *Galleria mellonella* and a virulent strain of *Staphylococcus aureus*, we investigate the infection dynamics and treatment outcomes of antibiotics of different classes—including antibiotics to which the bacteria are resistant—and a lytic bacteriophage. Surprisingly, we find that the host’s ability to control bacterial density and survive infection does not depend on the specific type of antimicrobial agent nor the bacteria’s susceptibility to it. Our results demonstrate that the innate immune system is the primary factor in therapeutic success, capable of clearing even highly resistant infections, such as those involving beta-lactamase-producing or ribosomal mutant-resistant strains. These findings challenge the traditional antibiotic-centric view of infection outcomes and emphasize the need to account for host-pathogen-drug interactions beyond simple MIC measurements when designing clinical treatment regimens.

## Main Text

Contemporary antibiotic treatment is informed by the pharmacodynamics and pharmacokinetics of the treating antibiotics and infecting bacteria (1). Antibiotic pharmacodynamics (PD) considers two parameters: (i) the minimum concentration of the drug necessary to prevent the replication of the target bacteria (the MIC), and (ii) the relationship between the concentration of the drug and the rates of growth or death of the treated bacteria (2). Pharmacokinetics (PK) is the rate of change in the drug’s effective concentration in the treated patient. PK depends on the rates of adsorption, distribution, metabolism, inactivation, and excretion of the treating drug (2). During treatment, PD and PK are considered together across time, particularly the amount of time at which the concentration of the antibiotic exceeds the MIC of the target bacteria, or as the area under the curve of the concentration of the drug in the plasma over time. Beyond MIC and the relationship between the rate of growth/death of the bacteria in vitro, alternative PD functions and parameters, such as those presented in (3), have been poorly exploited in the dosing and timing of treatment of antibiotic therapy. Stated another way, MIC is often the sole PD parameter employed in clinical decisions to use or not use an antibiotic.

Most critically, contemporary antibiotic treatment regimens fail to account for the host’s innate and adaptive immune responses to bacterial infection and its treatment. Animal-based approaches have been proposed to address this failing, such as the murine thigh infection model (4). These models are frequently employed under unrealistic conditions that do not reflect common human infections: animals are often treated within hours of infection (rather than when they are sick), and they are often immunocompromised.

In clinical medicine, the “precautionary principle” has been frequently overapplied; instead of treating effectively for the specific infection, we treat any infection as if we were in the worst clinical scenario. However, previous studies have shown that various antibacterial agents, such as antibiotics traditionally described as strongly bactericidal, weakly bactericidal, or bacteriostatic and even bacteriophage (phage) can yield similar survival outcomes (5-7).

To investigate the dynamics of antimicrobial therapy in a realistic and experimentally tenable animal model, we use the larvae of the Wax Moth, *Galleria mellonella* (8). Previous studies have demonstrated that this system is amenable to studies of chemotherapeutic action, particularly in the role of the innate immune system in treatment outcome (5). We have recently shown that this system is exceptionally faithful at testing quantitative hypotheses in infection biology. Using these larvae and a highly virulent strain of *Staphylococcus aureus (9)*, we explore the treatment dynamics of a bacteriostatic and bactericidal drug to which the bacteria are sensitive, two bactericidal drugs to which the bacteria are resistant via varying mechanisms, and a highly lytic bacteriophage to which the bacteria are susceptible. Surprisingly, we find that the ability of the treated larvae to control the density of infecting bacteria does not depend on the type of treating agent nor the sensitivity of the bacteria to the treating drug. This unexpected result challenges the century-long antibiotic-centric view of the outcome of treated infected individuals, suggesting that the primary factor in therapeutic success is, in fact, the efficacy of the natural immunological response of the host.

## Results

### The dynamics of infection without treatment

We first consider the dynamics of infection without treatment and then determine the dynamics of infection with treatment. In Figure 1, we present data previously published in (5) to demonstrate the distributions that obtain with infections in the absence of treatment. Notably, the mean density of bacteria per larva by four hours is ∼10^5^ CFU regardless of whether the initial inoculum was high (10^8^ CFU), medium (10^6^ CFU), or low (10^4^ CFU). By 24 hours post-injection, while the average density of bacteria per larva for the low and medium inocula is still ∼10^5^ CFU per larva, for the high inoculum, the average has increased to 10^8^ CFU per larva. Notably, all larvae with an infection density above 10^6^ CFU per larva succumbed to the infection regardless of inoculum size. For morbidity and mortality data, see Supplemental Table 1.

**Figure 1.**
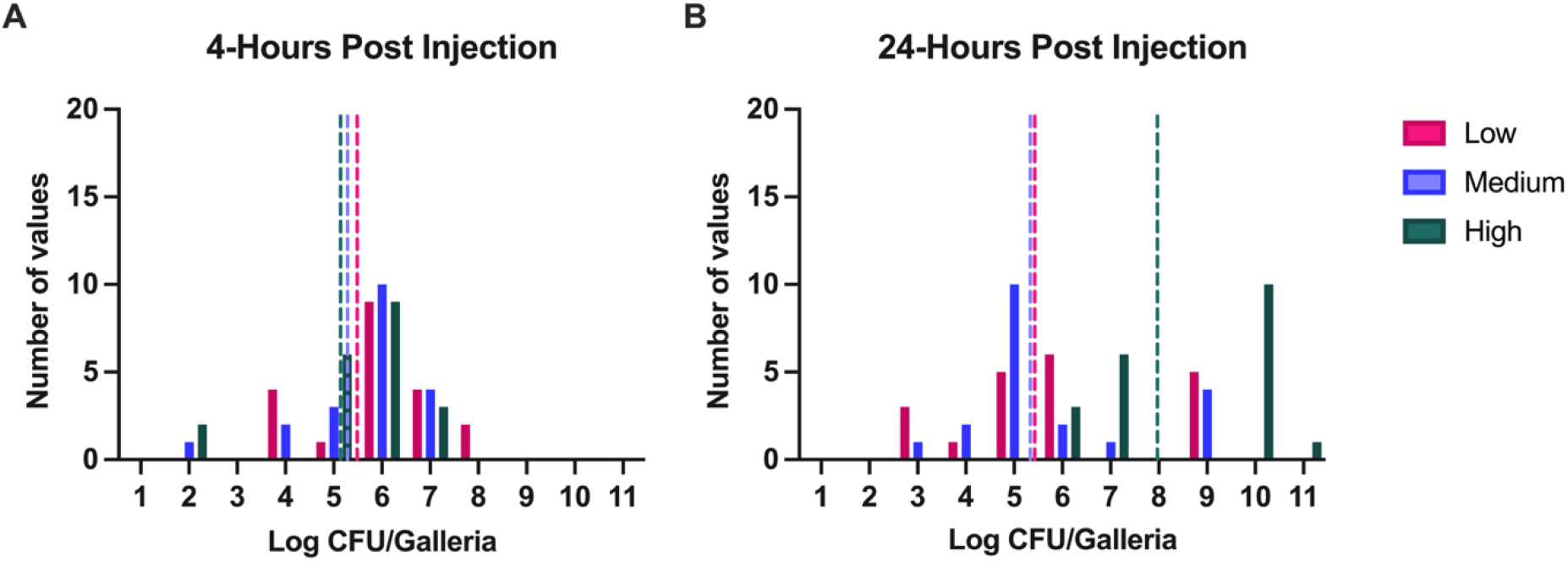
Dynamics of infection in the absence of treatment. Larvae were infected with three different densities of *S. aureus* MN8: low (10^4^ CFU/larva; pink), medium (10^6^ CFU/larva; purple), or high (10^8^ CFU/larva; green) and were sacrificed after (A) 4 or (B) 24 hours post-infection. Shown are the number of larvae within each log_10_ CFU. The mean of the CFU at each time point for each inoculum density is presented as a dotted line of the same color. For each condition, N=20 individual larvae were inoculated.

### The dynamics of treatment with an antibiotic to which the bacteria are susceptible

In humans, there is no difference in survival when either bactericidal or bacteriostatic drugs are used to treat bacterial infections. Our previous experiments have confirmed that both survival and the degree of suppression of the bacterial infection in *Galleria* do not differ between these categories of antibiotics. To determine whether the dynamics of treatment with these agents differ, we use daptomycin (a highly bactericidal drug; Figure 2) and linezolid (a bacteriostatic drug against *Staphylococcus*; Figure 3). By E-test and time kill curves, *S. aureus* MN8 would be classified as susceptible to both drugs (Supplemental Table 2; Supplemental Figure 1). For both drugs, we investigate treated larvae in three conditions: treated immediately and sacrificed at 4 hours (A); treated immediately and sacrificed at 24 hours (B); and treated only once phenotypic signs of infections begin to show and then sacrificed at 24 hours (C). In terms of survival, these agents do not differ at 4 h, and there are only minimal differences at 24 h. However, when treatment is delayed until sickness, linezolid performs substantially worse with a high degree of mortality (Supplemental Table 3 and 4). Regarding the dynamics of infections treated with these drugs, linezolid can suppress and even clear the infection more effectively than daptomycin, even with a low initial inoculum. With a medium and a high initial inoculum, both drugs can control the infection to nearly the same degree; however, the differences in terms of morbidity and mortality are stark at the high inoculum despite this lack of dynamic difference.

**Figure 2.**
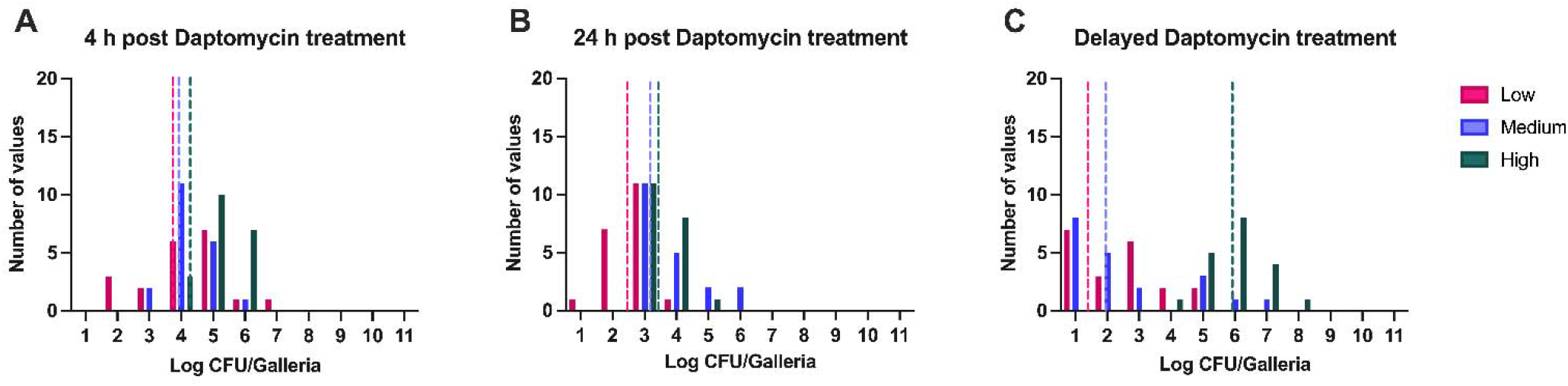
Dynamics of infection with daptomycin treatment. Larva were infected with three different densities of *S. aureus* MN8: low (10^4^ CFU/larva; pink), medium (10^6^ CFU/larva; purple), or high (10^8^ CFU/larva; green) and were sacrificed after 4 (A) or 24 (B and C) hours post-infection. Treatment occurred either immediately (A and B) or once symptoms of infection began to show (C). Shown are the number of larvae within each log_10_ CFU. The mean of the CFU at each timepoint for each inoculum density is presented as a dotted line of the same color. For each condition, N=20 individual larvae.

**Figure 3.**
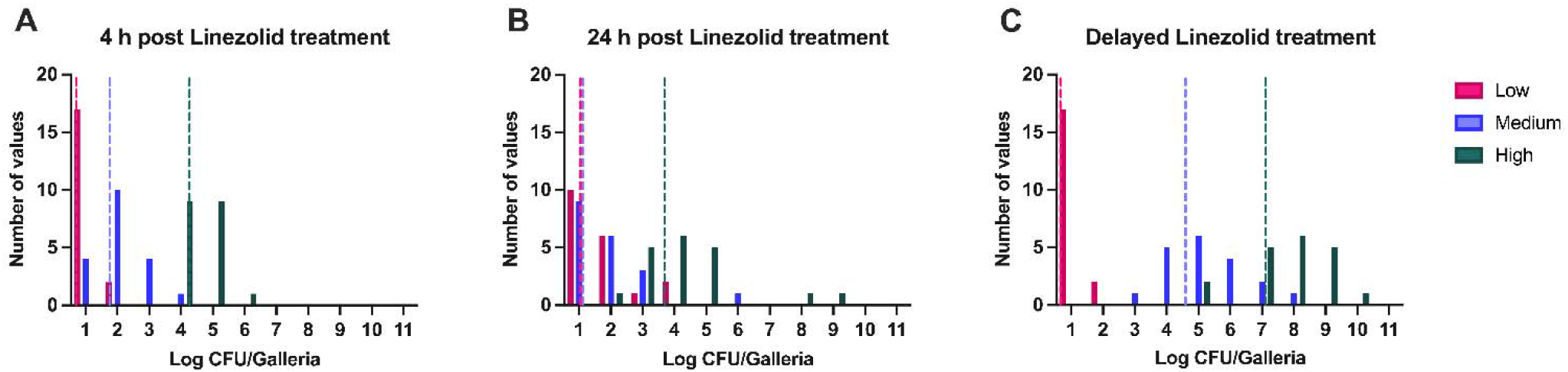
Dynamics of infection with linezolid treatment. Larvae were infected with three different densities of *S. aureus* MN8: low (10^4^ CFU/larva; pink), medium (10^6^ CFU/larva; purple), or high (10^8^ CFU/larva; green) and were sacrificed after 4 (A) or 24 (B and C) hours post-infection. Treatment occurred either immediately (A and B) or once symptoms of infection began to show (C). Shown are the number of larvae within each log_10_ CFU. The mean of the CFU at each time point for each inoculum density is presented as a dotted line of the same color. For each condition, N=20 individual larvae.

### The dynamics of treatment with an antibiotic to which the bacteria are resistant

The decision to use or not use an antibiotic is based almost solely on the susceptibility of the bacteria to that drug, most frequently determined exclusively by estimating the MIC of the drug in vitro. To ascertain the extent to which the dynamics of treatment differ due to antibiotic resistance, we use ampicillin (Figure 4) and streptomycin (Figure 5). By E-test, *S. aureus* MN8 would be classified as highly resistant to both drugs (Supplemental Table 2; Supplemental Figure 1), albeit with different mechanisms. The strain MN8 contains a beta-lactamase gene, *blaZ*, a penicillinase whose expression is controlled via beta-lactam induction and thus is resistant to ampicillin. For streptomycin, on the other hand, MN8 was marked via spontaneous mutagenesis, selecting for *rpsL* ribosomal mutants, allowing this strain to be distinguished from the larvae’s microbiome. This streptomycin resistance is extremely robust and stable. For both drugs, we investigate treated larvae in three conditions: treated immediately and sacrificed at 4 hours (A); treated immediately and sacrificed at 24 hours (B); and treated only once phenotypic signs of infections begin to show and then sacrificed at 24 hours (C). In terms of survival, both drugs ameliorate morbidity and mortality at four hours. However, at 24 hours, streptomycin does a better job at preventing mortality than ampicillin, while both drugs perform better than no treatment (Supplemental Table 5 and 6). When treatment is delayed, ampicillin can prevent sickness and death in all cases, whereas streptomycin cannot decrease mortality when the initial inoculum is high. In terms of bacterial dynamics, the drugs clearly control the infection in all cases—in fact, in many cases, these antibiotics lead to infection clearance, something not seen without treatment. Interestingly, the only difference between these two agents is at 4 hours with a low initial inoculum (Figure 4A). In this case, ampicillin cannot initially control the infection, but by 24 hours, it has done so.

**Figure 4.**
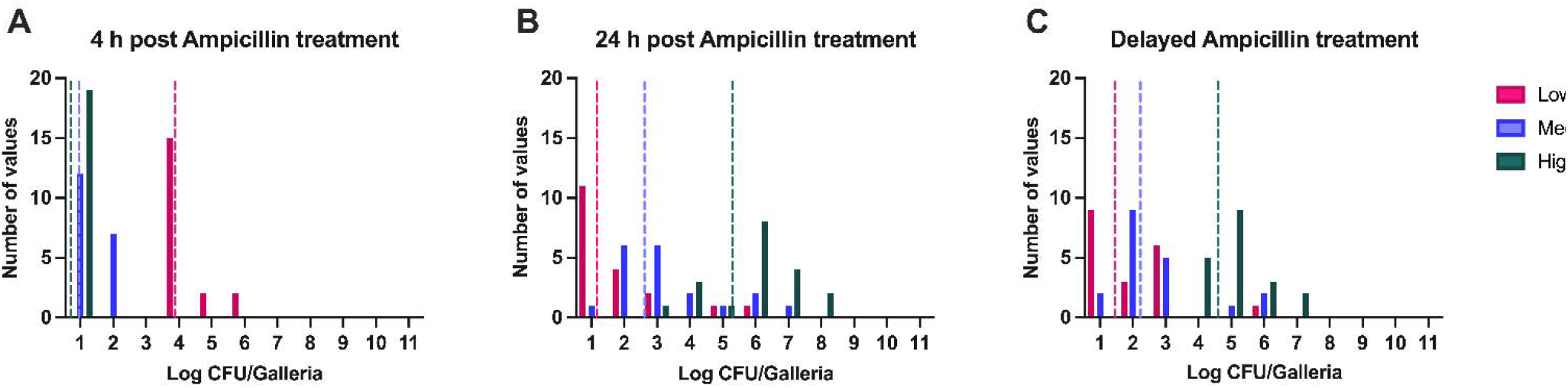
Dynamics of infection with ampicillin treatment. Larvae were infected with three different densities of ampicillin-resistant *S. aureus* MN8: low (10^4^ CFU/larva; pink), medium (10^6^ CFU/larva; purple), or high (10^8^ CFU/larva; green) and were sacrificed after 4 (A) or 24 (B and C) hours post-infection. Treatment occurred either immediately (A and B) or once symptoms of infection began to show (C). Shown are the number of larvae within each log_10_ CFU. The mean of the CFU at each time point for each inoculum density is presented as a dotted line of the same color. For each condition, N=20 individual larvae were inoculated.

**Figure 5.**
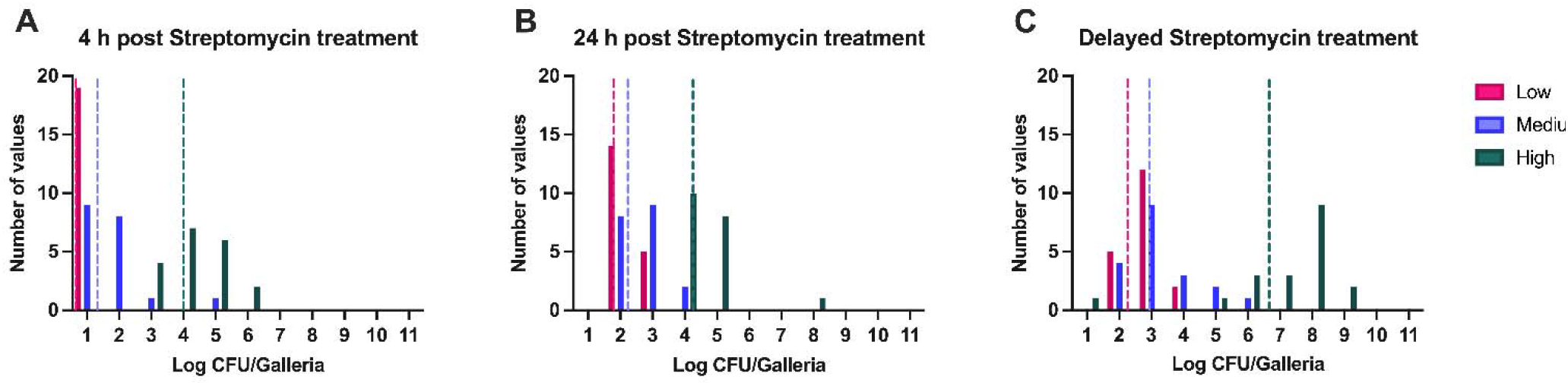
Dynamics of infection with streptomycin treatment. Larvae were infected with three different densities of a streptomycin-resistant mutant of *S. aureus* MN8: low (10^4^ CFU/larva; pink), medium (10^6^ CFU/larva; purple), or high (10^8^ CFU/larva; green) and were sacrificed after 4 (A) or 24 (B and C) hours post-infection. Treatment occurred either immediately (A and B) or once symptoms of the infection began to show (C). The number of larvae within each log_10_ CFU is shown. The mean of the CFU at each time point for each inoculum density is presented as a dotted line of the same color. For each condition, N=20 individual larvae were inoculated.

### The dynamics of treatment with bacteriophage

Given the increasing frequency of infections with antibiotic-resistant pathogens, there has been a resurgence of interest in using phages to treat bacterial infections. Here, we explore the dynamics of infection under treatment with a well-studied, highly virulent *S. aureus* phage, PYO^Sa^ (Figure 6) (10), in the same manner as above. Similar to antibiotics, the action of the phage requires growing bacteria. However, unlike antibiotics, which work in a drug-concentration-dependent manner, the action of phage depends on both the densities of the phage and the bacteria. In terms of mortality, the phage does an excellent job at preventing morbidity and mortality at low and medium initial bacterial inocula. With a high initial bacterial inoculum, the phage reduces morbidity and mortality but shows greater success when treatment is delayed (Supplemental Table 7). Regarding the dynamics, the phage does an excellent job at suppressing the bacterial infection in all cases. With a low inoculum, the phage clears the infection most of the time; however, with a high inoculum, these viruses only control the infection but do not clear it (Figure 6 A, B, and C). Regarding the phage density, there is a direct correlation between the recovered densities of the host bacteria and the phage, as expected.

**Figure 6.**
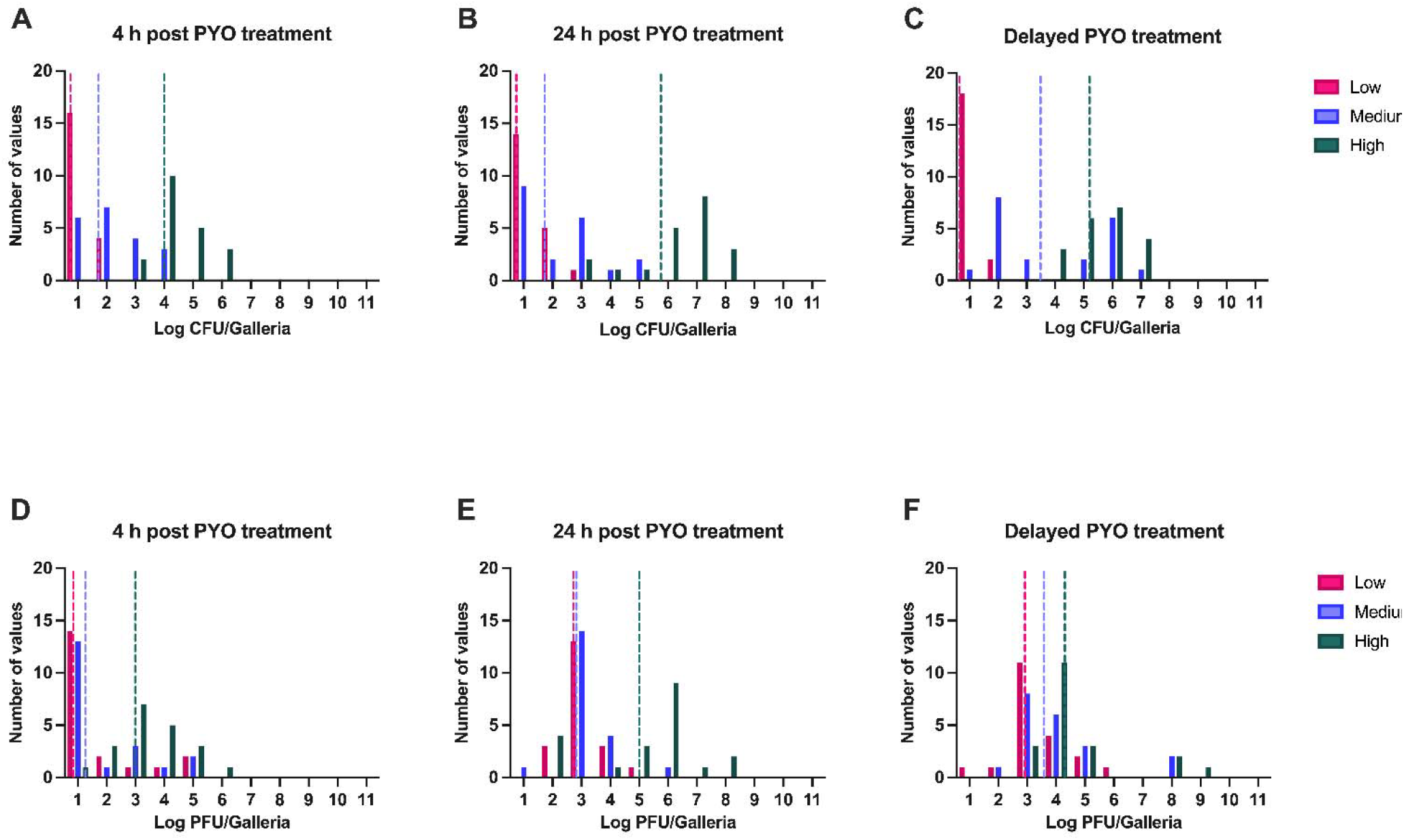
Dynamics of infection with phage treatment. Larvae were infected with three different densities of *S. aureus* MN8: low (10^4^ CFU/larva; pink), medium (10^6^ CFU/larva; purple), or high (10^8^ CFU/larva; green) and were sacrificed after 4 (A) or 24 (B and C) hours post-infection. Treatment occurred either immediately with 10^8^ PFU/larva (A and B) or once symptoms of infection began to show (C). Shown are the number of larvae within each log_10_ CFU (A through C) or the number of larvae within each log_10_ PFU (D through F) for each respective experiment. The mean of the CFU or PFU at each time point for each inoculum density is presented as a dotted line of the same color. For each condition, N=20 individual larvae were inoculated per condition.

## Discussion

For almost a century, antibiotics have gained the reputation of being the most powerful agents in combatting bacterial infections (11, 12). Consequently, bacterial susceptibility or resistance to these drugs, measured in most cases by the minimal inhibitory concentration (MIC), has been considered the primary critical factor to determine which antibiotic should be used in treatment and to predict its therapeutic success. This measure is often singularly employed epidemiologically to monitor the spread of antibiotic resistance rather than a clinical outcome such as treatment failure (13).

Given the antibiotic resistance crisis and the culture of fear surrounding the failure of antibiotics, an image has been presented of apocalyptic landscapes arising from antibiotic resistance. To reduce antibiotic resistance, antibiotic policies and surveillance of the emergence and spread of resistance in different pathogens have been universally implemented (14). These policies have led to widely applied antibiotic-use restriction protocols (15). The real burden of antibiotic resistance, particularly the mortality it causes, is challenging to evaluate due to a multiplicity of confounding factors (16, 17). Frequently, the rate of consumption of antibiotics correlates with resistance only in low-income countries, as the lack of sanitation and the poor nutritional conditions reinforces the spread of resistant organisms (18).

We should revise our antibiotic-centric view of the treatment of infections. The infective process occurs in a changing and complex landscape; additionally, most infections where antibiotics are used would be self-limited due to the contribution of the innate immune response (19). In most common infections, such as otitis media in children, antibiotics play only a marginal role in restoring health *(20)*.

Our results demonstrate the utility of the *Galleria* model in studying the treatment outcomes and dynamics of antibiotic therapy in infected individuals. We predict that an immunodeficient *Galleria* should provide similar results to conventional antibiograms regarding antibiotic resistance, leading to treatment failure. Indeed, we expect that the results obtained with the *Galleria* model would also be obtained in more traditional models, such as mice. Importantly, experiments in *Galleria* have been shown to be highly reproducible, and this model significantly reduces the cost and workload associated with animal experiments while increasing the replication (21-23). Our results support the use of this system for the study of single antimicrobial chemotherapeutics and open the door to investigating the joint application of these agents, such as the co-administration of bacteriophage and antibiotics.

The resistance dynamics observed for ampicillin and streptomycin further underscore the dominant role of the host immune system over purely pharmacological parameters. In the case of ampicillin, the resistance mechanism of *S. aureus* MN8 is driven by the production of beta-lactamase, which creates a density-dependent clearance problem where individual bacteria can only neutralize so much drug. Despite this high level of resistance, many larvae remained healthy following treatment, suggesting that ampicillin itself was not the primary driver of the infection clearance. Instead, this outcome highlights the critical contribution of the innate immune response, which effectively controls the infection when the bacterial burden is within a manageable threshold. Mortality was primarily observed in larvae receiving a high initial inoculum where the immune response is naturally less effective due to the overwhelming bacterial density. Similar dynamics were observed with streptomycin, where larvae infected with highly resistant mutants survived in most cases following therapy, except if the initial inoculum was excessively high. In these instances, the innate immune response does successfully control the resistant population.

Remarkably, the results for both ampicillin and streptomycin treatment did not differ substantially from those treated with antibiotics to which the bacteria were fully susceptible, such as daptomycin or linezolid. These findings collectively suggest that while resistance mechanisms, such as beta-lactamase production or ribosomal mutations can be seen *in vitro* via an increase in MIC, their impact on the ultimate trajectory of an infection *in vivo* is heavily modulated by the host’s immunological response.

Taken together, our study demonstrates that drugs traditionally described as bacteriostatic and bactericidal antibiotics and bacteriophages similarly lead to the clearance of infecting bacteria and ameliorate morbidity and mortality; and that, antibiotic-resistant bacteria can also be readily cleared with the antibiotic to which the bacteria are resistant. These results emphasize the key role of the innate immune response in the control of infections and the need to consider factors other than MIC at the time of prescribing.

## Materials and Methods

### Growth media

All experiments were conducted in Muller Hinton II (MHII) Broth (90922-500G) obtained from Millipore. All bacterial quantification was done on Lysogeny Broth (LB) agar (244510) plates obtained from BD. E-tests were performed on MH agar plates made from MH broth (M391-500g) with 1.6% agar obtained from HiMedia.

### Growth and infection conditions

All experiments were conducted at 37 °C.

### Bacterial strains

All experiments were performed with *S. aureus* MN8 obtained from Tim Read of Emory University. *S. aureus* MN8 was marked with streptomycin resistance to enable differential plating from the larva microbiota.

### Antibiotics

Daptomycin (D2446) was obtained from Sigma-Aldrich. Streptomycin (S62000) was obtained from Research Products International. Ampicillin (A9518-25G) was obtained from Sigma-Aldrich. Linezolid (A3605500-25g) was obtained from AmBeed. All E-test strips were obtained from Biomérieux.

### Bacteriophage

The bacteriophage PYO^Sa^ was obtained from the Levin Laboratory’s bacteriophage collection.

### Bacteriophage preparation for injection

Lysates of PYO^Sa^ were grown on *S. aureus* Newman such that the total volume exceeded 250 mL of media. These initial lysates were spun down and filtered through a 0.22 μm filter to remove cellular debris. From there, the lysates were run through a 100 kD tangent flow filtration (TFF) cassette (PAL 0A100C12) with a PAL Minimate TFF system. The volume was reduced during this process to 15 mL which was refiltered through a 0.22 μm filter. This lysate was incubated with shaking at 25 °C with equal parts octanol (Thermo Scientific 4345810000) for 24 hours. The non-organic fraction was then dialyzed with Thermo Scientific’s 250 kD float-a-lyzer cassette (66455) against 60% ethanol and then again against saline. Finally, the lysate was refiltered through a 0.22 μm filter.

### Sampling bacterial and phage densities

Bacteria and phage densities were estimated by serial dilutions in 0.85% NaCl solution followed by plating. The total density of bacteria was estimated on LB (1.6%) agar plates. To estimate the densities of free phage, chloroform was added to suspensions before serial dilutions. These suspensions were mixed with 0.1 mL of overnight MHII grown cultures of wild-type *S. aureus* MN8 in 4 mL of LB soft (0.65%) agar and poured onto semihard (1%) LB agar plates.

### *G. mellonella* preparation

*G. mellonella* larvae were obtained from Premium Crickets (Georgia, USA) and placed immediately at 4 °C for 48 hours as a cold shock. Larvae were then sorted such that only those that weighed between 180 and 260 mg.

### *G. mellonella* infection and sampling

Larvae were allowed to acclimatize at 37 °C for 48 hours before the experiment. Overnight cultures of *S. aureus* MN8 were centrifuged for 20 min at room temperature at maximum speed, the supernatant discarded, and the pellet resuspended in saline. This process was repeated 5 times. The bacterial suspension was diluted in saline to the desired inoculum concentration. Groups of 10 larvae were injected with 5 μL of the desired bacterial suspension at the last left proleg. Larvae were then treated either immediately or once sickness began to show by injection in the last right proleg. Larvae were incubated at 37 °C in the dark without food. After 4 or 24 hours the larvae were placed in 1 mL of saline and then homogenized. The homogenate was plated in LB agar plates containing 400 μg/mL of streptomycin.

### *G. mellonella* treatment

Larva were treated by injection in the last right proleg. Antibiotic concentrations were analogous to those used for clinical treatment in humans and the amount of antibiotic determined by weight for a total final treatment amount of: LIN-0.002 mg per larvae; TET-0.01 mg per larvae; DAP-0.002 mg per larvae; AMP-0.04 mg per larvae; STR-0.003 mg per larvae. Phage was delivered in the same way with a fixed concentration of 10^8^ PFU per larvae.

### Antibiotic minimum inhibitory concentration

Susceptibility to each treating antibiotic was determined by E-test on MH agar plates.

### Statistical analysis

All statistical analyses were performed in GraphPad Prism Version 8.0.1 using a two-tailed unpaired parametric t-test.

## Supporting information

All Supplemental Materials

## Acknowledgments

We would like to thank Alysha Ismail for her excellent laboratory management work and assistance with the *in vivo* bacteriophage experiment as well as the rest of the Levin Lab for the support.

## Funding Support

We thank the U.S. National Institute of General Medical Sciences for their funding support via R35 GM 136407 and the U.S. National Institute of Allergy and Infectious Diseases for their funding support via U19 AI 158080 to BRL. FB acknowledges the support of CIBERESP (CB06/02/0053) from the Carlos III Institute of Health of Spain. The funding sources had no role in the design of this study and will not have any role during its execution, analysis, interpretation of the data, or drafting of this report. The content is solely the responsibility of the authors and do not necessarily represent the official views of the National Institutes of Health nor those of the Carlos III Institute of Health of Spain.

## Data Availability

All data underlying this report can be found in the main text and its figures or the supplemental materials. Access to bacterial strains and phages can be had by contacting the corresponding author.

## Notes

### Competing Interest Statement

The authors have declared no competing interest.

