## Supplementary material for "The role of the innate immune system in shaping the dynamics of antimicrobial treatment": All Supplemental Materials

**This file includes:**

Supplemental Figure 1

Supplemental Tables 1 to 7

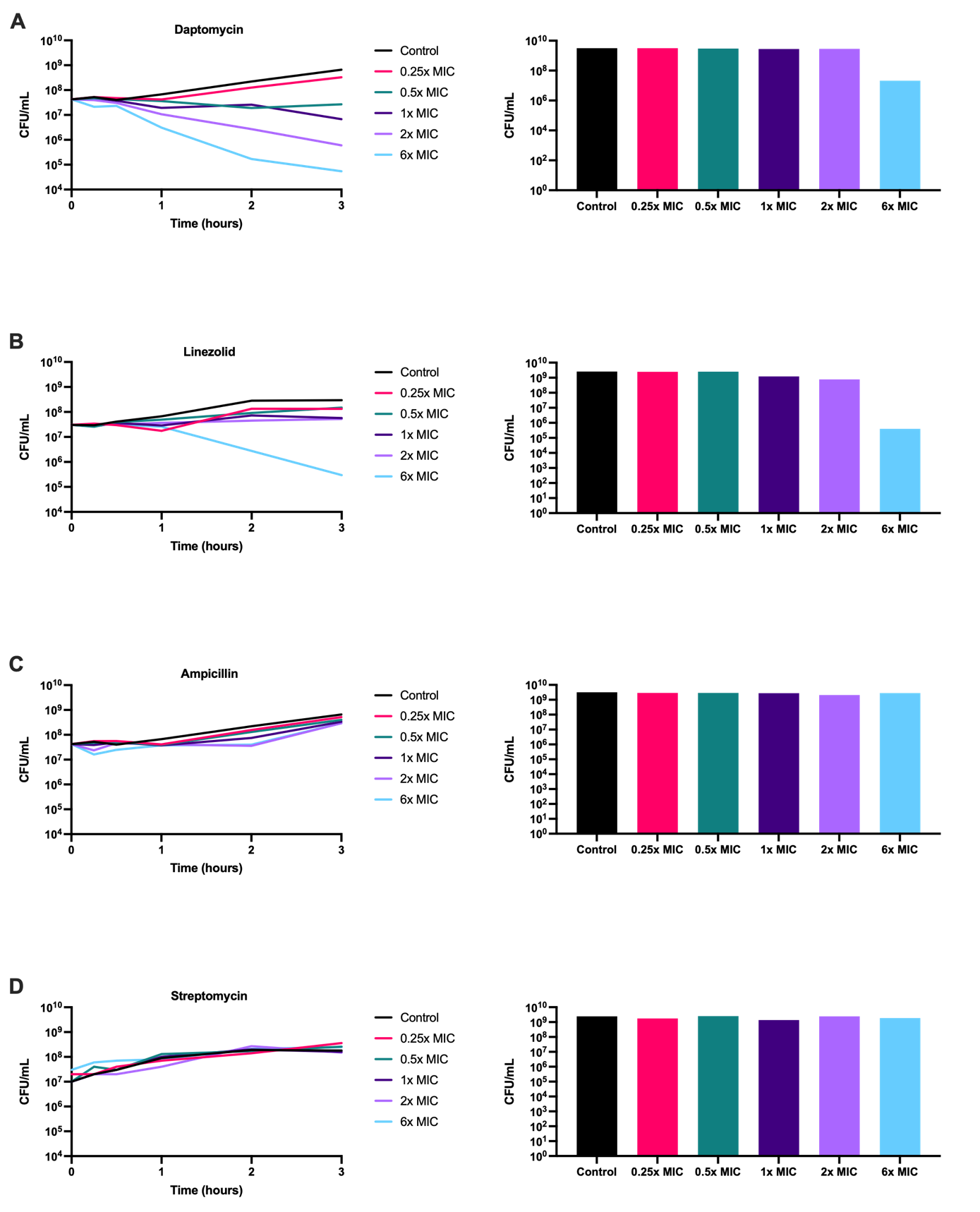

**Supplemental Figure 1. Time Kill Curves of MN8 and the Antibiotics used in Treatment of the *Galleria*.** MN8 was exposed to either 0x MIC (black) 0.25x MIC (red), 0.5x MIC (green), 1x MIC (dark purple), 2x MIC (light purple), 6x MIC (blue) of daptomycin (A), Linezolid (B), Ampicillin (C), or streptomycin (D). Presented on the left is the change in bacterial density over three hours. The density of bacteria at 24 h is presented in the right column.

**Supplemental Table 1. Infected Larvae Without Treatment**

| **Treatment** | **Inoculum** | **Healthy** | **Sick** | **Dead** |
| --- | --- | --- | --- | --- |
| **Kill at 4 Hours** | Low | 19 | 1 | 0 |
|  | Medium | 17 | 2 | 1 |
|  | High | 18 | 2 | 0 |
| **Kill at 24 Hours** | Low | 15 | 0 | 5 |
|  | Medium | 16 | 0 | 4 |
|  | High | 3 | 4 | 13 |

**Supplemental Table 2**. **MIC of *Staphylococcus aureus* MN8 to Treating Antibiotics**

| **Antibiotic** | **MIC via E-Test (µg per mL)** |
| --- | --- |
| Daptomycin | 0.5 |
| Linezolid | 0.5 |
| Ampicillin | >1024 |
| Streptomycin | >4096 |

**Supplemental Table 3. Infected Larvae Treated with Daptomycin**

| **Treatment** | **Inoculum** | **Healthy** | **Sick** | **Dead** |
| --- | --- | --- | --- | --- |
| **Immediate Kill at 4 Hours** | Low | 20 | 0 | 0 |
|  | Medium | 20 | 0 | 0 |
|  | High | 14 | 6 | 0 |
| **Immediate Kill at 24 Hours** | Low | 20 | 0 | 0 |
|  | Medium | 20 | 0 | 0 |
|  | High | 17 | 3 | 0 |
| **Delayed Treatment Kill at 24 Hours** | Low | 17 | 1 | 2 |
|  | Medium | 15 | 1 | 4 |
|  | High | 9 | 7 | 4 |

**Supplemental Table 4. Infected Larvae Treated with Linezolid**

| **Treatment** | **Inoculum** | **Healthy** | **Sick** | **Dead** |
| --- | --- | --- | --- | --- |
| **Immediate Kill at 4 Hours** | Low | 20 | 0 | 0 |
|  | Medium | 20 | 0 | 0 |
|  | High | 18 | 2 | 0 |
| **Immediate Kill at 24 Hours** | Low | 20 | 0 | 0 |
|  | Medium | 20 | 0 | 0 |
|  | High | 16 | 2 | 2 |
| **Delayed Treatment Kill at 24 Hours** | Low | 20 | 0 | 0 |
|  | Medium | 17 | 2 | 1 |
|  | High | 3 | 1 | 16 |

**Supplemental Table 5. Infected Larvae Treated with Ampicillin**

| **Treatment** | **Inoculum** | **Healthy** | **Sick** | **Dead** |
| --- | --- | --- | --- | --- |
| **Immediate Kill at 4 Hours** | Low | 20 | 0 | 0 |
|  | Medium | 20 | 0 | 0 |
|  | High | 20 | 0 | 0 |
| **Immediate Kill at 24 Hours** | Low | 19 | 1 | 0 |
|  | Medium | 19 | 1 | 0 |
|  | High | 7 | 7 | 6 |
| **Delayed Treatment Kill at 24 Hours** | Low | 20 | 0 | 0 |
|  | Medium | 20 | 0 | 0 |
|  | High | 16 | 3 | 1 |

**Supplemental Table 6. Infected Larvae Treated with Streptomycin**

| **Treatment** | **Inoculum** | **Healthy** | **Sick** | **Dead** |
| --- | --- | --- | --- | --- |
| **Immediate Kill at 4 Hours** | Low | 20 | 0 | 0 |
|  | Medium | 20 | 0 | 0 |
|  | High | 20 | 0 | 0 |
| **Immediate Kill at 24 Hours** | Low | 19 | 1 | 0 |
|  | Medium | 20 | 0 | 0 |
|  | High | 7 | 12 | 1 |
| **Delayed Treatment Kill at 24 Hours** | Low | 19 | 0 | 1 |
|  | Medium | 19 | 1 | 0 |
|  | High | 6 | 3 | 11 |

**Supplemental Table 7. Infected Larvae Treated with Bacteriophage**

| **Treatment** | **Inoculum** | **Healthy** | **Sick** | **Dead** |
| --- | --- | --- | --- | --- |
| **Immediate Kill at 4 Hours** | Low | 20 | 0 | 0 |
|  | Medium | 20 | 0 | 0 |
|  | High | 20 | 0 | 0 |
| **Immediate Kill at 24 Hours** | Low | 20 | 0 | 0 |
|  | Medium | 20 | 0 | 0 |
|  | High | 7 | 6 | 7 |
| **Delayed Treatment Kill at 24 Hours** | Low | 20 | 0 | 0 |
|  | Medium | 18 | 1 | 1 |
|  | High | 14 | 2 | 4 |
